# Fast transcriptional activation of developmental signalling pathways during wound healing of the calcareous sponge *Sycon ciliatum*

**DOI:** 10.1101/2021.07.22.453456

**Authors:** Cüneyt Caglar, Alexander Ereskovsky, Mary Laplante, Daria Tokina, Sven Leininger, Ilya Borisenko, Genevieve Aisbett, Di Pan, Marcin Adamski, Maja Adamska

**Affiliations:** Research School of Biology, Australian National University, Canberra, Australia; ARC Centre of Excellence for Coral Reef Studies; Institut Méditerranéen de Biodiversité et d’Ecologie Marine et Continentale (IMBE), CNRS, Marseille, France; Saint-Petersburg State University, Saint-Petersburg, Russia; Sars International Centre for Marine Molecular Biology, University of Bergen, Bergen, Norway

## Abstract

**Background:** The ability to regenerate lost or damaged body parts is an ancient animal characteristic with a wide yet variable distribution across all phyla. Sponges, likely the sister group to all other animals, have remarkable regenerative abilities including whole body regeneration and re-development from dissociated cells. The calcareous sponge *Sycon ciliatum* has been subject to various regeneration studies since the beginning of the last century. However, the early steps of wound healing of *S. ciliatum* have not been addressed from the molecular perspective.

**Results:** In this study, we combined electron microscopy with gene expression analysis to investigate wound healing after transverse sectioning of *S. ciliatum*. Microscopic analysis revealed massive transdifferentiation and collective migration behaviour of choanocytes and pinacocytes early upon injury (6-12h) as the main mechanisms for quick closure of the wound surface. RNA-sequencing identified upregulation of components of the conserved metazoan Wnt and TGFβ signalling pathways within 3h, preceding morphologically detectable wound healing events. De novo upregulation after a decline in expression coincides with morphologically visible polarity establishment. Moreover, by integrating the new wound healing data set with previously published data derived from intact sponge, we demonstrate similarity between gene activity during early wound healing and osculum maintenance. Whole mount *in situ* hybridisation of the TGFβ signalling pathway ligand SciTGFβU and signal transducer SciSmadRa show that the early activation of both is initially encompassing a large area surrounding the cut surface with gradual restriction to the edge of the forming regenerative membrane as wound healing progresses. While SciTGFβU transcripts are localised to exo- and endopinacocytes, SciSmadRa expression appears across all cell types. Using an EdU cell proliferation assay, we found that a global increase in cell proliferation is not visible before 12h into wound healing. Hence, the initial stages to cover the injury site including cell transdifferentiation and migration seem to be executed by cells remaining after injury. Gene expression clustering coupled with GO term enrichment analysis confirmed that expression of genes involved in processes related to cell proliferation, DNA repair as well as apoptotic processes at 3 and 6h of wound healing was not upregulated. On the other hand, genes associated with positive regulation of transcription, signal transduction, actin filament and chromatin organisation, as well as the Wnt signalling pathway are upregulated at early wound healing stages.

**Conclusion:** We have analysed wound healing in the calcareous sponge *Sycon ciliatum* using microscopic and genomic methods. This study highlights a remarkable mechanism of interplay between cell transdifferentiation and collective migration we hypothesise to be regulated by conserved metazoan developmental pathways and numerous taxonomically restricted genes. Expression of these genes in regenerating and intact sponges sheds light on the long-standing question whether embryonic developmental pathways are redeployed in regeneration.

## Background

Regeneration is an ancient and widespread trait in Metazoa, yet highly variable across phyla. An evolutionary developmental approach has led to a continuously growing body of knowledge on the underlying mechanisms and their conservation across animals, with an increasing number of diverse model species supporting these efforts [1, 2, 3, 4, 5, 6].

Sponges are likely the earliest branching animal lineage and are thus crucial to our understanding of the evolutionary origin of key animal traits [7, 8, 9]. They have a morphologically simple body plan lacking nerves, muscles and gut, instead consisting of two layers of epithelial cells and varying amount of non-epithelial tissue [10, 11, 12]. The epithelia are composed of two major cell types: pinacocytes, flattened cells, forming the outermost layer (exopinacoderm), as well as channels and atrium (endopinacoderm) in some sponge lineages, and flagellated choanocytes forming internal chambers and establishing a continuous water flow to facilitate feeding and oxygenation. The non-epithelial layer, mesohyl, is sandwiched between pinacoderm and choanoderm, and contains diverse cell types including gametes, skeleton-producing cells and amoeboid cells of multiple functions. Sponges possess exceptional regenerative capacities, early studies particularly emphasising their ability to restore lost body parts and even regenerate from fragmented or dissociated tissues (whole-body regeneration) [13, 14, 15, 16]. Recent studies provide a thorough description of regenerative processes in a variety of sponge species and highlight the exceptional morphological plasticity manifested in a combination of cell differentiation as well as transdifferentiation and increased cell motility as underlying mechanisms [17, 18, 19, 20, 21, 22]. Furthermore, studies of sponge regeneration have revealed complex interplays of cell dedifferentiation, proliferation and redifferentiation and added to our understanding of their peculiar regenerative morphogenesis and renewed appreciation of the independent evolution of different sponge lineages over a long geological time [17, 22].

Regeneration research in sponges has largely been focused on demosponges, by far the most speciose class of the phylum Porifera [24]. While calcareous sponges have been subject of extensive developmental studies over the last century and have provided valuable insights into early animal evolution, only few studies on regeneration in Calcarea are available – mostly restricted to morphological and histological observations [12]. Syconoid sponges in particular represent unique candidates for comparative regenerative studies, given their radial symmetry around a directional, apical to basal, axis and extensive developmental regulatory gene toolkit [25, 26, 27, 28]. Although sponges from the genus *Sycon* have long been known for their remarkable regeneration abilities, only recent research has provided transcriptomic insight into restoration from dissociated cells including comparison to postembryonic development in *S. ciliatum* [29], a calcareous sponge found along the coasts of Atlantic Europe [28, 30, 31]. However, *S. ciliatum* whole body or structural regeneration has not been previously studied using genomic and modern microscopic tools.

The induction of sophisticated developmental programmes for regeneration is dependent on efficient injury response and wound healing [32, 33, 34]. We distinguish between wound healing (early phase) and rebuilding of missing structures (later phases). In this study we set out to provide a comprehensive account on *S. ciliatum* wound healing. We used SEM to capture cellular events during wound healing and regeneration – from injury through finely orchestrated cellular events in the early stages of healing leading to efficient tissue remodelling. SEM observations were complemented by RNA-seq data, supported by integration of previously published intact sponge data [27]. Transcriptomic dynamics during wound healing were investigated including differentially expressed gene analysis and Gene Ontology (GO) enrichment analysis of gene clusters. We have further visualised cell proliferation patterns throughout wound healing and regeneration to support our SEM analysis. Finally, we revealed expression patterns of key components of the TGFβ signalling pathway with cell-level resolution. Our findings contribute to the overarching endeavour to establish syconoid sponges as models for animal regeneration, with their unique phylogenetic position as well as bauplan and genetic characteristics offering exciting insights to better understand the evolution of metazoan regeneration.

## Results

### *Sycon ciliatum* whole-body regeneration

The body of *S. ciliatum* has a cylindrical or fusiform shape narrowing towards the apical and basal extremities (Fig. 1A & B). The base of the sponge forms a massive peduncle, while the top bears a single exhalant opening, the osculum, surrounded by a crown of diactin spicules. The organisation of the aquiferous system is syconoid with choanocyte-lined radial chambers surrounding the endopinacocyte-lined atrial cavity.

**Figure 1.**
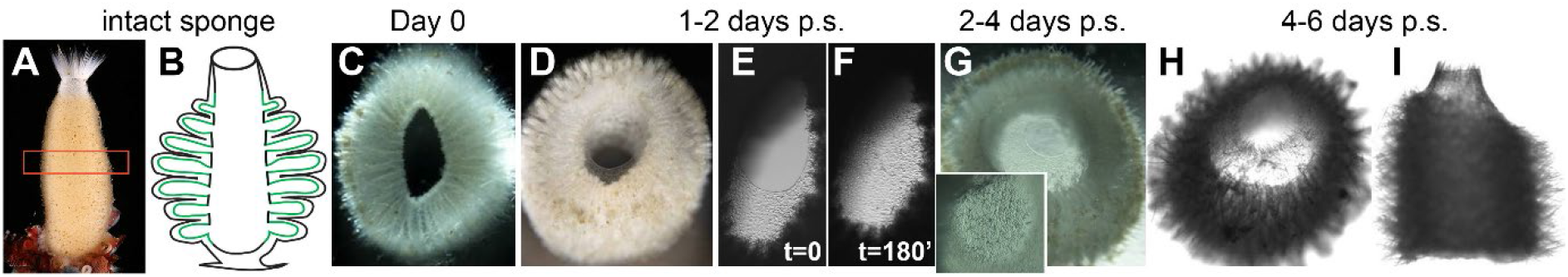
Sycon ciliatum regeneration time course. Intact sponges (A-B) were cut to obtain thin sections of the body column, which were maintained in Petri dishes with seawater (C). Between 1-2 days, a membrane is forming which is closing the atrium from the edges towards the centre (D-F). After 2-4 days, regenerative membrane has formed and a newly developing osculum is visible on the apical side (G). Regeneration is completed after 4-6 days with a clearly visible new osculum (H-I).

To gain initial insights into the regenerative capabilities of *S. ciliatum*, sections of its body column were dissected and maintained in Petri dishes with seawater (Fig. 1C). Apical and basal sides of obtained rings of sponge tissue are morphologically indistinguishable immediately after dissection. After 1-2 days, regenerative membrane (basal and apical membrane that covers the atrium during regeneration) formation can be observed, growing centripetally to cover the injury site (Fig. 1D–F). Once morphologically discernible, basal regenerative membrane formation is completed within a few hours (Fig. 1E & F). 2-4 days post sectioning, the regenerative membrane has fully covered the basal wound surface, while oscular opening is visible on the opposite end (Fig. 1G). Osculum formation, including a ring of diactines and the contractile sphincter, is completed after 4-6 days, which has been deemed completed regeneration (Fig. 1H & I).

### High cell plasticity and mobility during wound healing

To develop a deeper understanding of cellular mechanisms involved in *S. ciliatum* wound healing and regeneration, samples for electron microscopy were taken at 3, 6, 12, 24 hours, as well as 4 and 5 days post sectioning. Sponges were dissected perpendicularly along the apical-basal axis yielding rings of tissue with two injury sites except the most distal (osculum and basal) parts, which only have one injury site per ring. The regeneration can be roughly divided into four stages of characteristic morphological development. In particular, the early wound healing phase is characterised by dynamic and coordinated cellular processes to effectively cover the injury sites.

#### Stage 1: Wound healing 0 – 24 hours

Within the first three hours after cutting, the wound surface is covered with debris, consisting of microbes, fragments of cells and broken spicules (Fig. 2I). Choanocytes and some of the endopinacocytes have large phagosomes, which are absent in intact sponges. The shape of intact exopinacocytes surrounding the wound changes from flat to oval (Fig. 2J). Some intact, separated choanocytes at the wound surface as well as choanocytes at the zone surrounding the wound begin transdifferentiation. This is manifested in a cell shape change from trapeziform to oval and microvilli retraction (Fig. 2J–K). At the same time, intact endopinacocytes, that were situated proximal to the wound surface and were not damaged, begin to migrate towards the wound.

**Figure 2.**
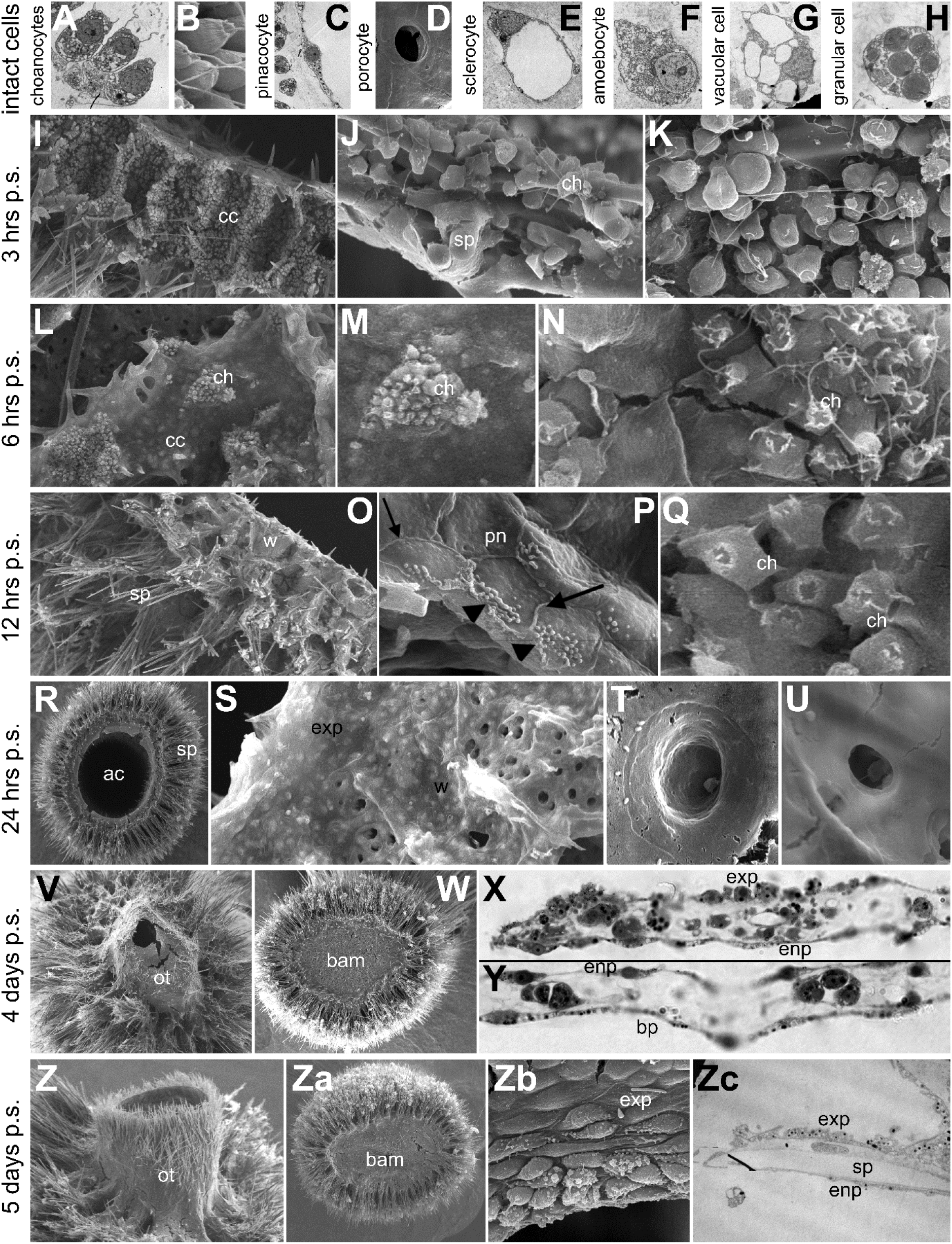
Scanning and transmission electron microscopy imagery of S. ciliatum cell types and regeneration time course from 3 hours to 5 days post dissection. Major cell types of S. ciliatum in their intact state for reference (A-H). Open choanocyte chambers and spicule fragments at the injury site and beginning of cellular events for wound healing between 3-6h (I-N). Major cell movement and transdifferentiation events of choanocytes and pinacocytes to restore functional epithelial layers at 12-24h (O-U). Regeneration is completed when development of the oscular tube is completed and the basal regenerative membrane fully covers the basal injury site, while cells establish normal shape around 4-5d after injury (V-Zc). Abbreviations: ac – atrial cavity, bam – basal regenerative membrane, cc – choanocyte chamber, ch – choanocyte, enp – endopinacocyte, exp – exopinacocyte, ot – oscular tube, pn – pinacocyte, sp – spicule, w – wound surface.

At approximately 6h after dissection, cleaning of the wound surface from debris and spicule fragments is largely finished, with only small zones of debris remaining. In the case that only a small part of a choanocyte chamber was dissected, the wound is covered with endopinacoderm and forms small islands of choanoderm (Fig. 2L & M). Big dissected choanocyte chambers remain opened at the wound surface and the choanocytes in these chambers maintain intact shape.

The choanocytes in marginal zones continue dedifferentiation and transdifferentiation into exopinacocytes: their microvilli are retracted while the cell’s basal part is flattened and merged with the pinacoderm, but flagella are still present (Fig. 2N). Undestroyed endopinacocytes that are on the wound surface follow their expansion by flattening and transdifferentiation into new exopinacoderm. The peripheral cells of this layer establish contacts (fuse) with intact choanoderm.

Between 6 and 12h, separated cells and cell layers display increased motility. All cells in and around the wound area show very active movement, dedifferentiation and transdifferentiation.

All (big and small) opened choanocyte chambers begin to be covered by new exopinacoderm (Fig. 2O). Migrating and flattening pinacocytes manifest a typical leading edge with pseudopodia in direction of the movement (Fig. 2P), while pinacocytes completely covering the wound do not feature any active movement. As a result of this collective cell migration, the wound is covered by a pinacoderm layer with stable intercellular junctions (Fig. 2P). Peripheral choanocytes at the margins of dissected choanocyte chambers continue their transdifferentiation into exo- and/or endopinacocytes (Fig. 2Q). However, groups of choanocytes or fragments of damaged choanocyte chambers can immerge in the mesohyl and form small spherical choanocyte chambers.

#### Stage 2: Atrial membrane and ostia development

The next phase of regeneration is characterised by the onset of regenerative membrane development (Fig. 2R). At 24h after dissection, the wound surface is covered with new exopinacoderm (Fig. 2S). The cells in this layer are very active, as manifested by the number of pseudopodia on their surface. This new epithelium forms ostia (incurrent pores) by transdifferentiation of some of the exopinacocytes into porocytes (Fig. 2T & U). A small invagination forms at the centre of the exopinacocyte, gradually deepening until breaking through and forming a cylindrical canal connecting the exopinacoderm with the underlying choanoderm. Newly growing exopinacoderm and intact endopinacoderm joint together at the internal border zone surrounding the atrium. Here, they form the regenerative membrane, a wrinkled bilayered epithelial structure (Fig. 2R). In the big dissected choanocyte chambers, which are not yet covered with new pinacoderm, the marginal choanocytes continue to transdifferentiate into pinacocytes. Some dedifferentiated, separated choanocytes migrate into the mesohyl and keep their typical basal flagellar apparatus (basal body and accessory centriole). No apparent cell proliferation or coordinated cell death events can be observed until this point.

After one day of regeneration, the first manifestations of axial polarity in different rings of the *Sycon* body can be observed. The most distal sponge body sections have only one injury surface, apical or basal, and therefore, these sections develop the corresponding regenerative membrane and restore the lost body parts. Dissected sections from the middle of the body column develop the apical and basal regenerative membranes in similar rates. Conservation of axial polarity is evident in these sections as they proceed to regenerate oscula at their apical part and basal structures (basopinacoderm including specialised skeleton) at the basal part.

#### Stage 3: Oscular tube and basal membrane with specialised skeleton

At 4 days of regeneration, an oscular tube has developed from the apical regenerative membrane and the basal regenerative membrane is completely closed (Fig. 2V & W). Both structures form specialised skeleton, characteristic for each of the parts. Basally external cells, basopinacocytes, actively secret “glue” for attachment to the substratum. Both, exo- and endopinacocytes are more active in the central zone of the developed oscular tube and passive at the peripheral surface. In the mesohyl of growing oscular membrane and basal regenerative membrane, cell concentration is increased, including amoebocytes, granular cells, and sclerocytes (Fig. 2X & Y).

#### Stage 4: Completed regeneration

At this final stage of regeneration, small sponges have formed from each of the body rings (Fig. 2Z & Za). While exo- and endopinacocytes of the oscular tube appear to be passive, there is active synthesis of new oscular and perioscular skeletal elements characteristic for this species (Fig. 2Z). In the apical part of the oscular tube a contractile sphincter is developing, which closes the cavity of the tube and regulates the water current (Fig. 2Zb). The sphincter has a double-layered structure and consists of exo- and endopinacoderm. The cells of the sphincter are very active in both layers, with exopinacocytes secreting glycocalyx at their external surface. The basal part of each new sponge is completely covered with basal regenerative membrane (Fig. 2Za). In the mesohyl of this layer (between endo- and basopinacoderm) are amoebocytes, sclerocytes, granular cells, and symbiotic bacteria (Fig. 2Zc).

### Cell proliferation is a late event in *S. ciliatum* regeneration

To investigate whether cell proliferation is associated with wound healing and regeneration, we utilised an EdU cell proliferation assay, applying the EdU one hour prior to fixation at different time points from 3 hours to 5 days (Fig. 3). Intact control sponges were kept in Petri dishes for the duration of the experiment to mitigate potential cell proliferation differences due to environmental conditions (Fig. 3A–B). Overall, intensity of fluorescence corresponds to rate of EdU incorporation, and distribution of labelled nuclei was compared in regenerating and intact sponges. The major proliferative population of cells are choanocytes, with no proliferation of pinacocytes observed in intact sponges or during regeneration. Interestingly, increased cell proliferation was not observed until 12-24h after injury (Fig. 3C–E), consistent with EM observations and suggesting that wound healing is solely accomplished by transdifferentiation of choanocytes and migration of pinacocytes. Only few labelled choanocytes could be recognised at 3h, with gradual increase from 12-24h stages, and dramatic increase visible at 48h and later stages. No remarkable cell proliferation could be observed in the developing membranes, which are mainly composed of pinacocytes (Fig. 3G). Perhaps more surprisingly, the increased intensity and distribution of DNA synthesis in cells during regeneration appears global (uniform across sections which are 3-4 radial chambers thick, Fig. 3H–I) rather than limited to the direct vicinity of the injury site. Once cell proliferation is initiated, the rate is similar in cells scattered throughout the entire tissue section from 48h to completion of regeneration. The late initiation and steady rate of cell proliferation until later stages of regeneration may underscore the importance of cell transdifferentiation and migration in wound healing. It appears that *S. ciliatum* has evolved a very efficient and quick wound healing mechanism in which cell proliferation is needed to replace recruited cells after the wound is closed.

**Figure 3.**
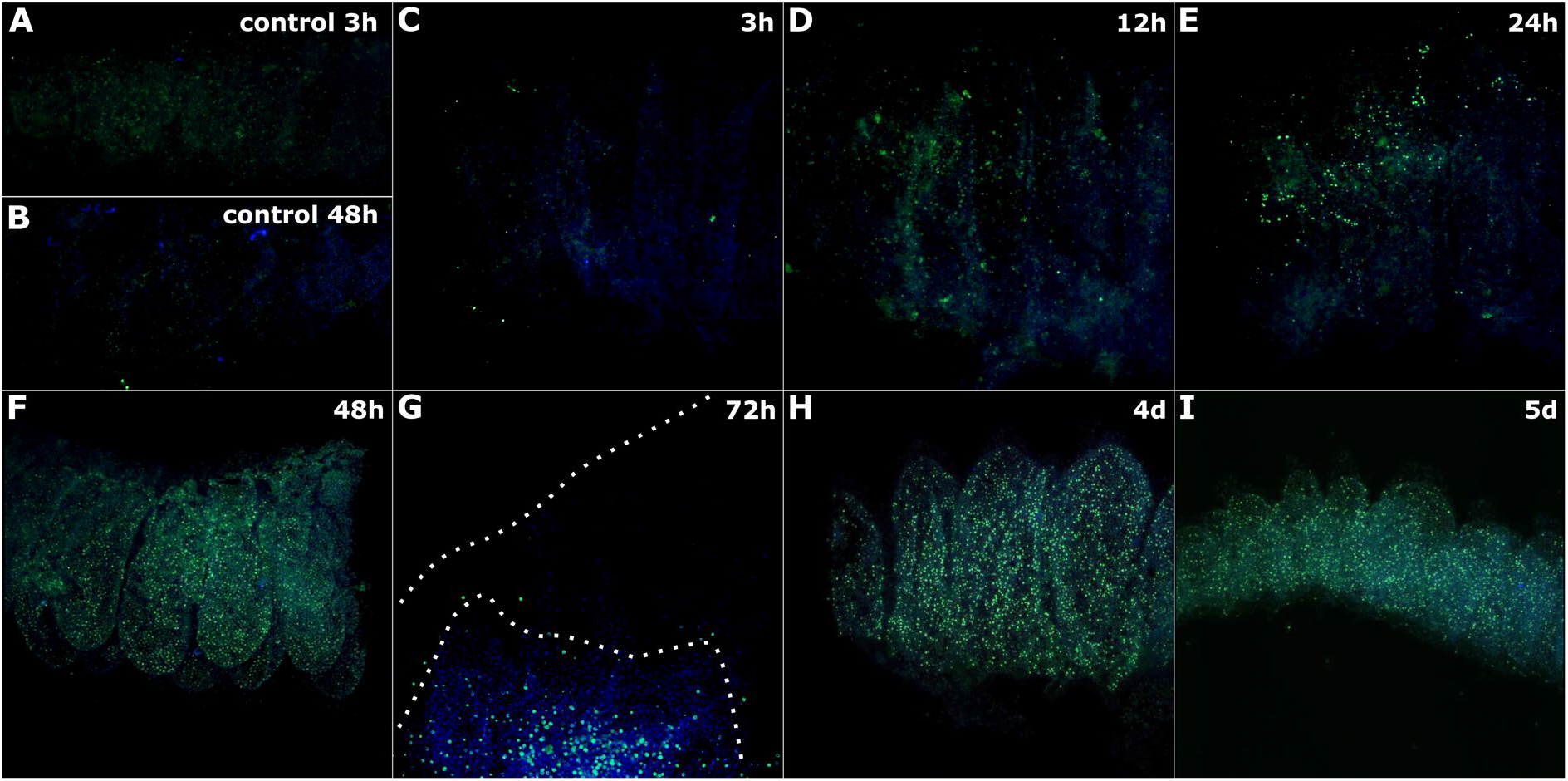
EdU proliferation assay of regenerating S. ciliatum transverse body sections. EdU was added 1 hour prior to fixation of the samples at indicated time points and was visualised through Alexa Fluor 488 detection (green). Nuclei are co-stained with DAPI (blue). White dotted line outlines the edges of the newly developing membrane at 72h. Sections A-D are shown from the cut surface, with atrium towards the top; in E tips of the radial chambers are pointing towards the viewer.

### Early wound healing and osculum maintenance show similarity in the transcriptome signatures

The morphological observations indicate that wound healing in *S. ciliatum* is a complex and dynamic process at the cellular level followed by a finely coordinated remodelling phase. We sought to identify genes involved in the wound healing stage of regeneration. It has been shown before that in intact sponges, the apical (oscular) region is characterised by specific expression of numerous developmental regulatory genes, homologues of which have pleiotropic roles in eumetazoans [27]. We thus wanted to find out if, and when, the osculum-associated genes become activated during regeneration. Therefore, this study incorporates datasets derived from two independent experiments conducted on different batches of sponges at different points in time. In addition to previously published gene expression data from axial slices of sponges processed immediately after sectioning (“intact sponge”, [27]), we sampled slices throughout the first 24h after sectioning (“wound healing”, this study) (Fig. 4A). As an internal control within the wound healing experiment, we sampled middle body sections of intact sponges that were kept under the same conditions as regenerating slices. To assess characteristics of the combined RNA-seq data, we performed principal component analysis on all available samples (Fig. 4).

**Figure 4.**
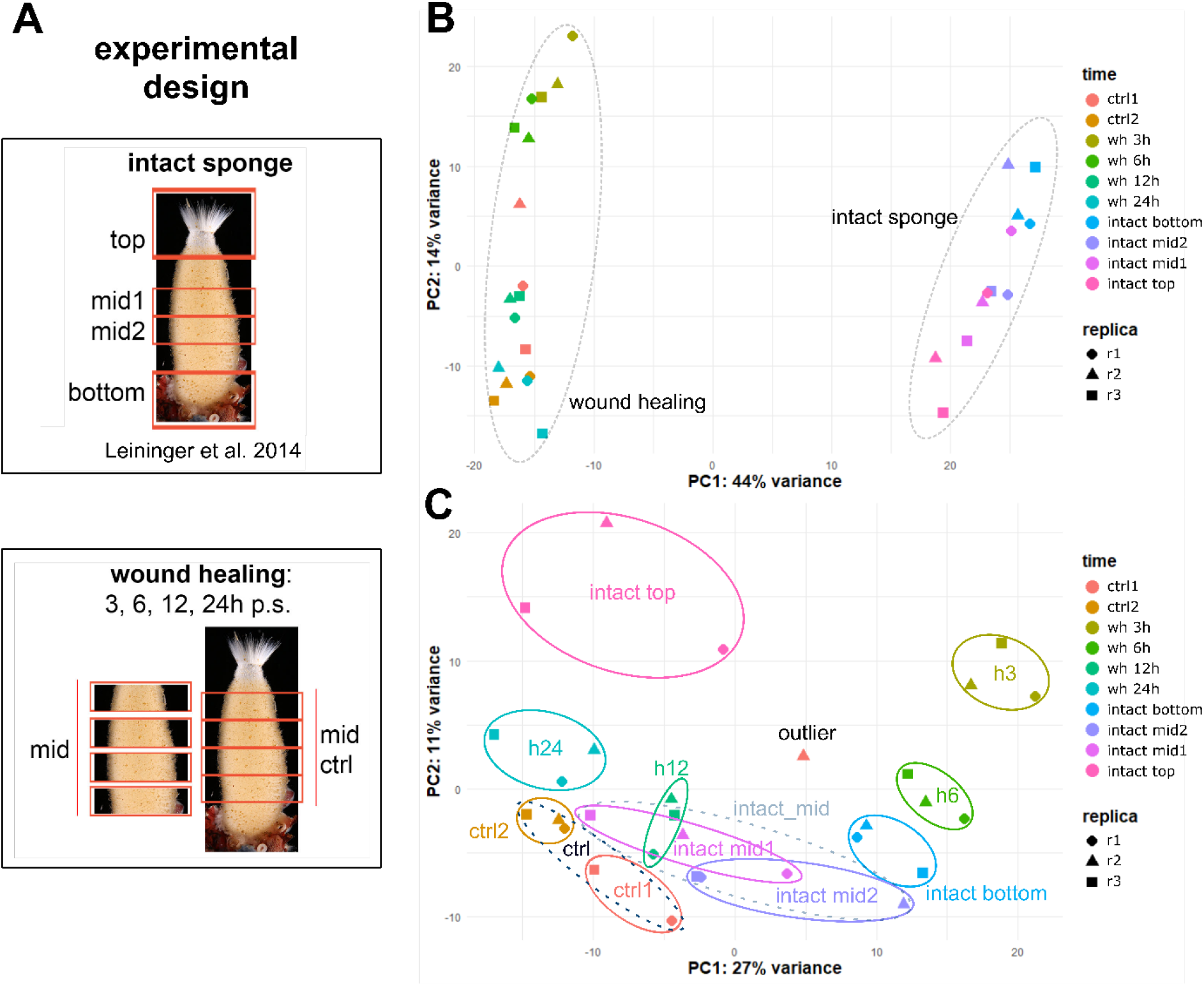
*Sycon ciliatum* gene expression: experimental design and principal component analysis of RNA-Seq data including intact sponge and wound healing experiments. This study utilises two transcriptomic experiments: 1) gene expression in different parts of an intact sponge and 2) wound healing (early regeneration) at 3-24 hours (**A**). Principal component analysis of intact sponge and wound healing RNA-Seq data before (**B**) and after (**C**) correction for batch effects. The PCA was performed using normalised RNA-Seq data of the top 500 most variable genes. After batch effect removal, samples from both experiments can be compared with each other.

The samples are separated by experiment in the first and second dimensions, which is expectedly caused by differences that occur in natural populations collected at different time points (batch effects). To integrate different RNA-seq datasets, batch effects need to be addressed prior to analysis which otherwise can lead to misleading conclusions [35, 36]. ComBat-seq was used to correct for batch effects and PCA was performed using batch adjusted and unadjusted data (Fig. 4B & C) [36]. Correction for batch-effects resulted in the samples of both experiments being intermixed in the first two dimensions, indicating that the batch effect removal was successful (Fig. 4C). In particular, the intact sponge mid-body sections and the control samples from the regeneration experiment cluster together (Fig. 4C, bottom left). With the exception for the second replicate of the 3-6h control in the wound healing data set (“outlier” in Fig. 5C), which was therefore excluded from further analysis. Tight clustering of replicates underlines good reproducibility of the data. Within PC1, the 3h wound healing samples are the farthest from all remaining samples, with the 6h samples positioned between 3h and the remaining samples (control, 12h and 24h; Fig. 4C). This result indicates that dramatic gene expression changes occur within the first 3 hours post-injury, and this initial response is then decreasing as the time progresses. Intriguingly, within PC2, the 3h regeneration time point shows highest similarity to the intact top (osculum) sample, followed by the samples collected 24h after injury. The similarity between the 24h stage and the osculum is not particularly surprising, given that new osculum formation begins at this stage of regeneration (Fig. 2R). However, the similarity of transcriptome profiles for 3h of wound healing and the intact top samples is remarkable. This is in line with our observations of increased cell mobility and transdifferentiation events at 3h, which is likely to utilise the complex repertoire of signalling genes making up the previously described specific genetic signature of *S. ciliatum* oscular tissue [27].

**Figure 5.**
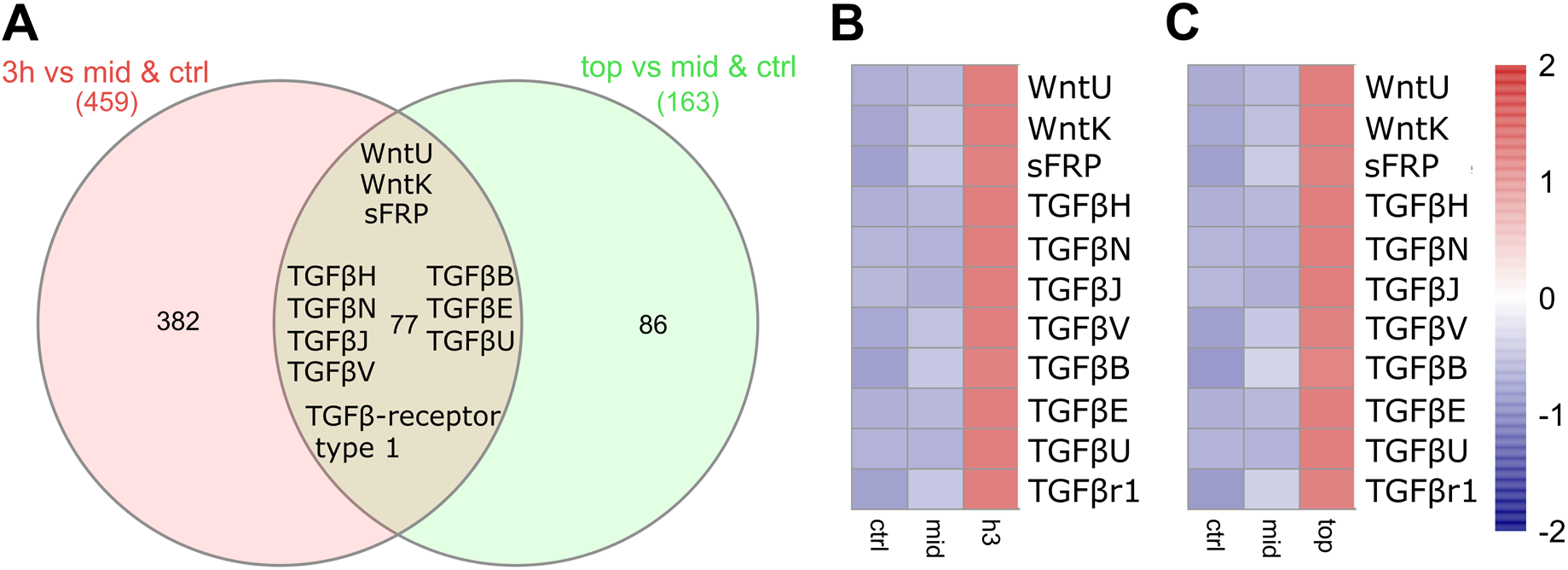
**A** Venn diagram illustrating the number of shared genes upregulated between control & intact mid vs h3h (red) and highly expressed between control & intact mid vs intact top (green) datasets. 77 genes are overlapping between both lists, which include developmentally important Wnt and TGFβ pathway components. Heatmaps showing expression of Wnt and TGFβ pathway genes at 3h (**B**) and top (**C**) compared to control and mid sponge section. Scaled heat map used to show their expression (z-score) based on summed count values of technical replicates.

### Early wound healing and osculum maintenance share developmental genes

A crucial question in developmental biology is the extent to which developmental toolkits are redeployed in wound healing and regeneration. To investigate whether cellular events in wound healing and osculum identity maintenance are controlled by similar genetic programmes, we performed pairwise comparisons between 3h and intact top samples, as well as each wound healing control and intact mid samples. First, we generated a stringent (at least two-fold enrichment and adjusted p-value ≤ 0.05) list of 77 genes which are upregulated at 3h and specifically enriched in the intact top by comparing both to control and intact mid and intersecting the resulting lists. Strikingly, among these 77 genes are 8 components of the TGFβ signalling pathway, and 3 components of the Wnt signalling pathway, both of which are known to have developmentally significant functions across eumetazoans (Fig. 5).

We have identified two Wnt and seven TGFβ ligands, a potential Wnt pathway modulator (sFRP) and one TGFβ receptor to be highly expressed at the earliest investigated wound healing stage (3h) and the osculum tissue of an intact sponge. This suggests an important role of the Wnt and TGFβ signalling pathways in wound healing and regeneration in addition to previously reported involvement in *S. ciliatum* development [27]. The remaining 66 genes are from diverse families involved in various biological processes, most notably, four of them are known to regulate actin dynamics and cell motility (supplementary material).

### Gene expression and GO term analysis of *S. ciliatum* wound healing

To investigate global gene expression trends and explore pathways and biological processes involved in *S. ciliatum* wound healing, we performed a broad analysis of the RNA-seq data. Firstly, genes with similar expression profiles were assigned to clusters using *Clust* [37]. Seven clusters of different expression patterns were generated, which in total include 13031 genes (Fig. 6A). Gene ontology (GO) enrichment analysis for biological process and molecular function in these clusters was conducted using *GOseq* [38]. Cluster C1 and C4 have a relatively high number of unique GO terms related to biological process and molecular function compared to other clusters, indicating major biological changes 3-6h after injury (Fig. 6A & B). Interestingly, genes upregulated at early wound healing stages are assigned to GO terms associated with positive regulation of transcription, signal transduction, actin filament and chromatin organisation, as well as the Wnt signalling pathway (Fig. 6A & C). Genes in cluster C4 have low expression at 3h and 6h and are involved in processes related to cell proliferation, DNA repair as well as apoptotic processes (Fig. 6A & D). This is in line with our morphological observations that indicate major developmental processes like transdifferentiation and cell migration in the absence of cell proliferation in the early stages of wound healing. While cell proliferation is essential for regeneration in many animal lineages, programmed cell death has been increasingly understood to induce and promote wound healing and regeneration [39, 40]. This is further evidence that cell proliferation and apoptotic processes are likely not driving the early stages of *S. ciliatum* regeneration, and that tissue remodelling appears to rely on collective migration of existing cells.

**Figure 6.**
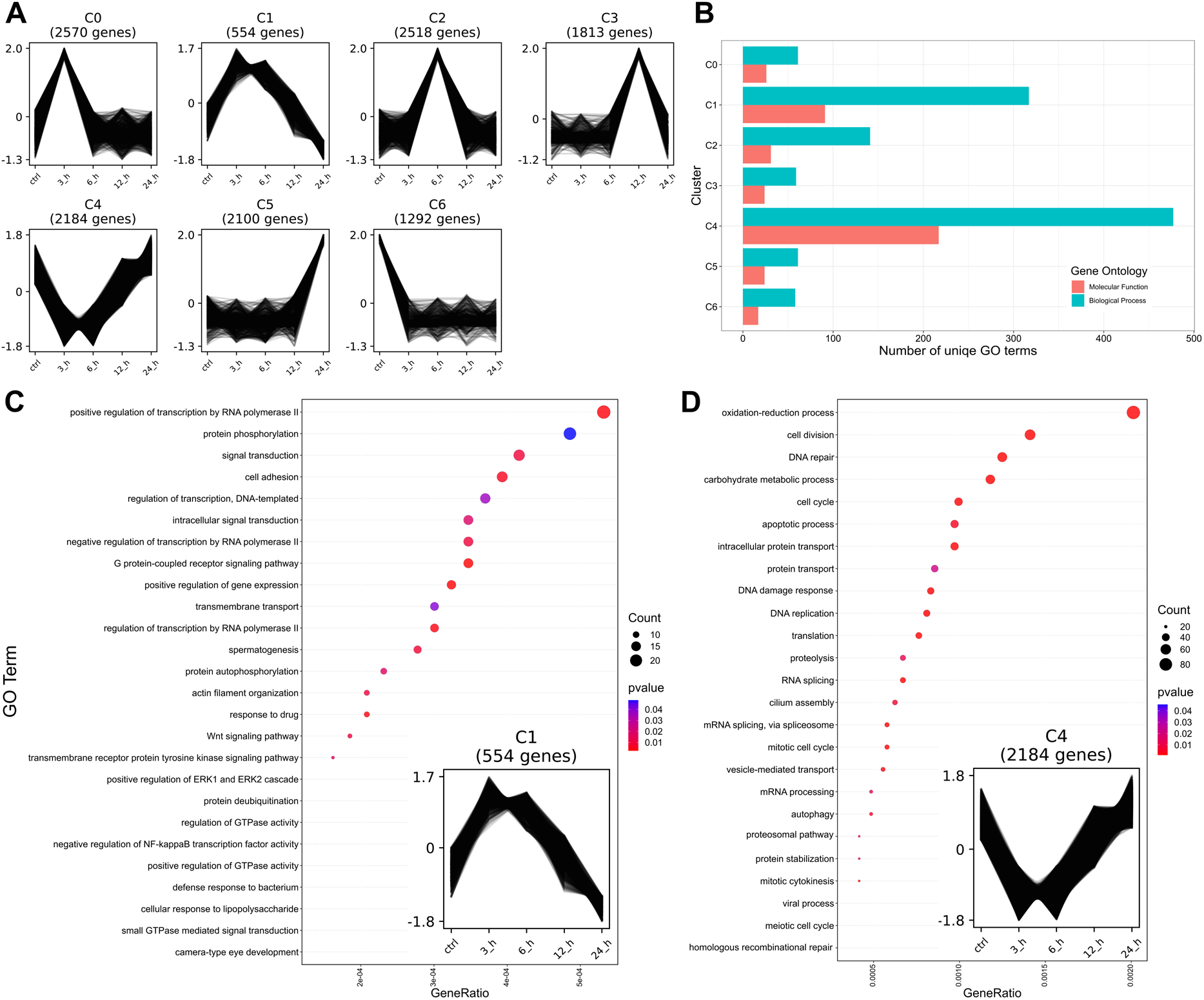
Transcriptional dynamics throughout *S. ciliatum* wound healing. **A** Clustering of 13031 genes into seven clusters according to their expression values. **B** Number of unique gene ontology (GO) terms related to molecular function and biological process by gene expression clusters (A). **C** GO term enrichment analysis of clusters C1 (554 genes) and C4 (2184) genes showing the top most enriched biological processes terms. For better visualisation a cut-off value of six (C1) and 16 (C4) were chosen showing the most enriched 26 and 25 biological processes, respectively.

To provide additional insights into global differential gene expression dynamics in response to injury, we conducted a time course analysis of wound healing using the R/Bioconductor package *timecourse* [41, 42]. All genes (83952 in total) were included in the analysis, which were first scanned for ranks of Wnt and TGFβ signalling pathway components. TGFβU was ranked highly among these genes (rank 108), highlighting a potentially essential role in *S. ciliatum* regeneration. We have also identified previously known bHLH transcription factors like SciSCLb (rank 5) and SciMyoRb (211), both expressed during embryonic development [43]. However, an overwhelming proportion of highly ranked genes (74 in top 100) are unknown or taxonomically restricted, underlining the fact that taxon specific genes may be crucial regulators of developmental processes in any investigated system. Function of the majority of these taxonomically restricted genes in *S. ciliatum* remains unknown. However, previous studies have identified a number of biomineralisation genes and demonstrated their temporal and spatial expression patterns during (carbonate) skeleton development in *S. ciliatum* [44, 45]. Two of these genes are involved in diactin (two rays) and triactin (three rays) spicule formation by specialised cells. SciDiactinin (rank 147) and SciTriactinin (rank 6094) are possibly involved in the restoration and reshaping of the new sponge individual after injury, as both are upregulated early during wound healing, preceding the initial morphological signs of spicule formation.

### Expression of Wnt and TGFβ pathway components in *S. ciliatum* wound healing

As the Wnt and TGFβ signalling pathways stand out as possible regulators of regeneration in *S. ciliatum*, we have analysed the expression profiles of individual components throughout the wound healing (Fig. 7). A group of genes including ligands, receptors, signal transducers and a transcription factor of the Wnt pathway is upregulated quickly in response to injury within 3h, while a larger group of Wnt pathway genes is upregulated at later stages of regeneration at 12-24h (Fig. 7A). An overwhelming proportion of ligands and signal transducers of the TGFβ pathway are highly upregulated within 3h upon injury, most decreasing expression steadily during wound healing, while some show a downward trend followed by a de novo upregulation at 24h (Fig. 7B).

**Figure 7.**
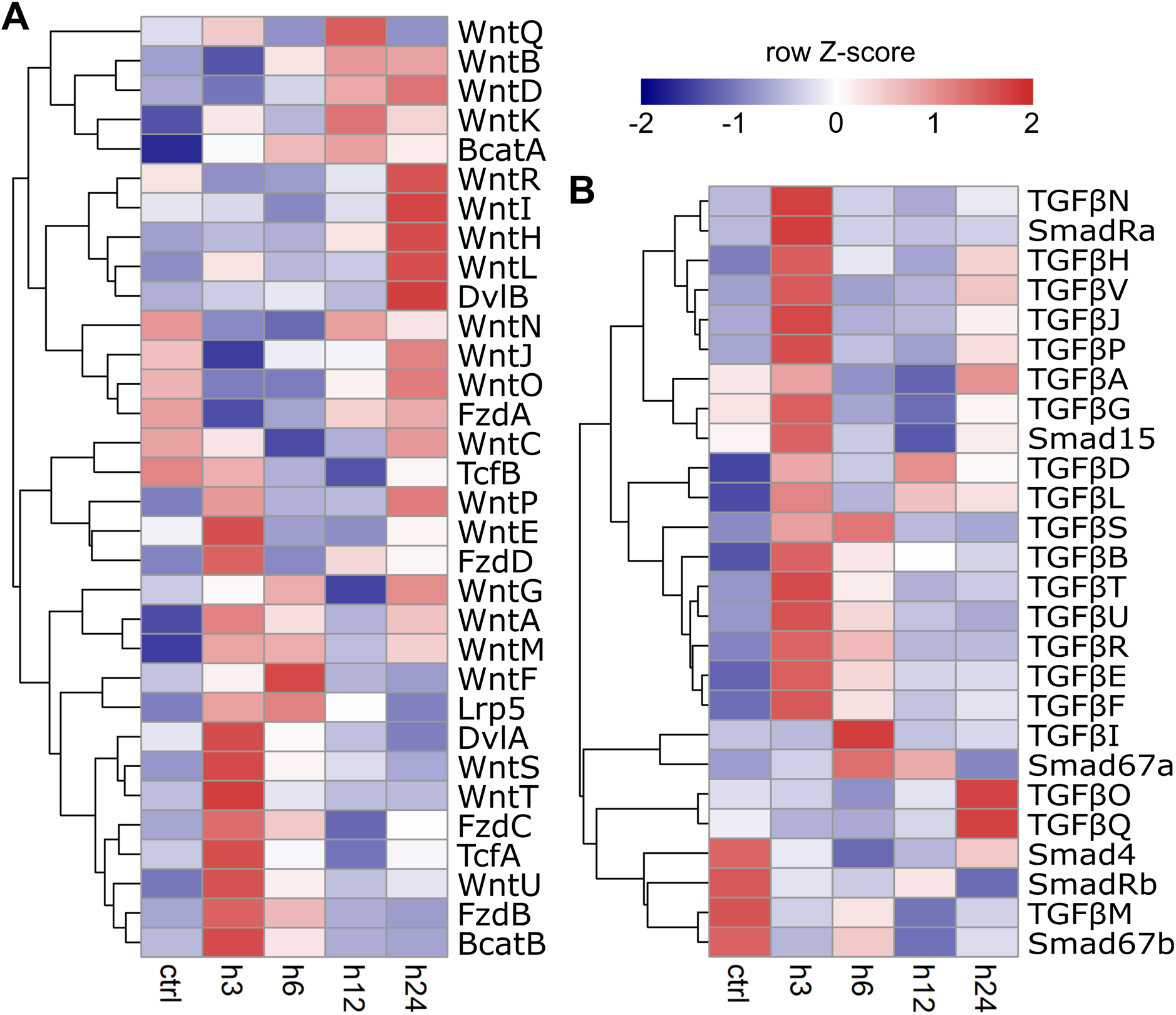
Scaled heat maps illustrating the expression (*z-score*) of previously described *S. ciliatum* (**A**) Wnt and (**B**) TGFβ pathway components throughout wound healing. Expression is based on summed count values of technical replicates.

This initial upregulation of Wnt and TGFβ pathway components suggests an involvement in transdifferentiation and/or cell migration events during early wound healing. Subsequent upregulation of Wnt pathway osculum marker genes clearly corresponds to the period when the polarity of the sponge sections becomes morphologically evident. These genes are likely involved in maintenance of the apical – basal polarity and re-establishment of corresponding structures during regeneration.

While bulk transcriptome analysis allows unbiased identification of genes and pathways involved in the investigated biological process, it lacks cell type and position resolution. We have therefore selected two genes combining high fold-change with high level of expression to carry *in situ* hybridisation analysis throughout the early regeneration stages: SciTGFβU and SciSmadRa (Fig. 8). Expression of one of the ligands of the TGFβ pathway, SciTGFβU is virtually absent from non-oscular somatic tissues ([27] and Fig. 8A). Transcriptome analysis indicated that expression of this gene dramatically (approximately 500-fold) increases by 3h post-injury, and then gradually decreases, but is still above control levels at 24h (Fig. 8A). Consistent with the transcriptome analysis, *in situ* hybridisation revealed strong expression of SciTGFβU at 3h post injury, with no detectable staining in the control samples (which were fixed before sectioning) (compare Fig. 8 B to C–E). The expression domain was gradually reduced from its observed peak at 3h, when it covers up to three radial chambers away from the cut (Fig. 8 C), till 48h when the positive cells could be observed only at the very edge of the forming regeneration membrane (Fig. 8D, E). The transcripts are localised to exopinacocytes (Fig. 8F) and endopinacocytes (Fig. 8G), with no other cell types appearing positive. Strikingly, while in intact sponges’ expression of this gene is limited to the osculum, in the regenerating fragments the basal and apical surfaces of the cut show indistinguishable expression levels (Fig. 8C, F, G). The expression of SciTGFβU is consistent with autocrine activation of the TGFβ pathway in migrating pinacocytes, in line with reports on key role of this pathway in collective cell migration [46].

**Figure 8.**
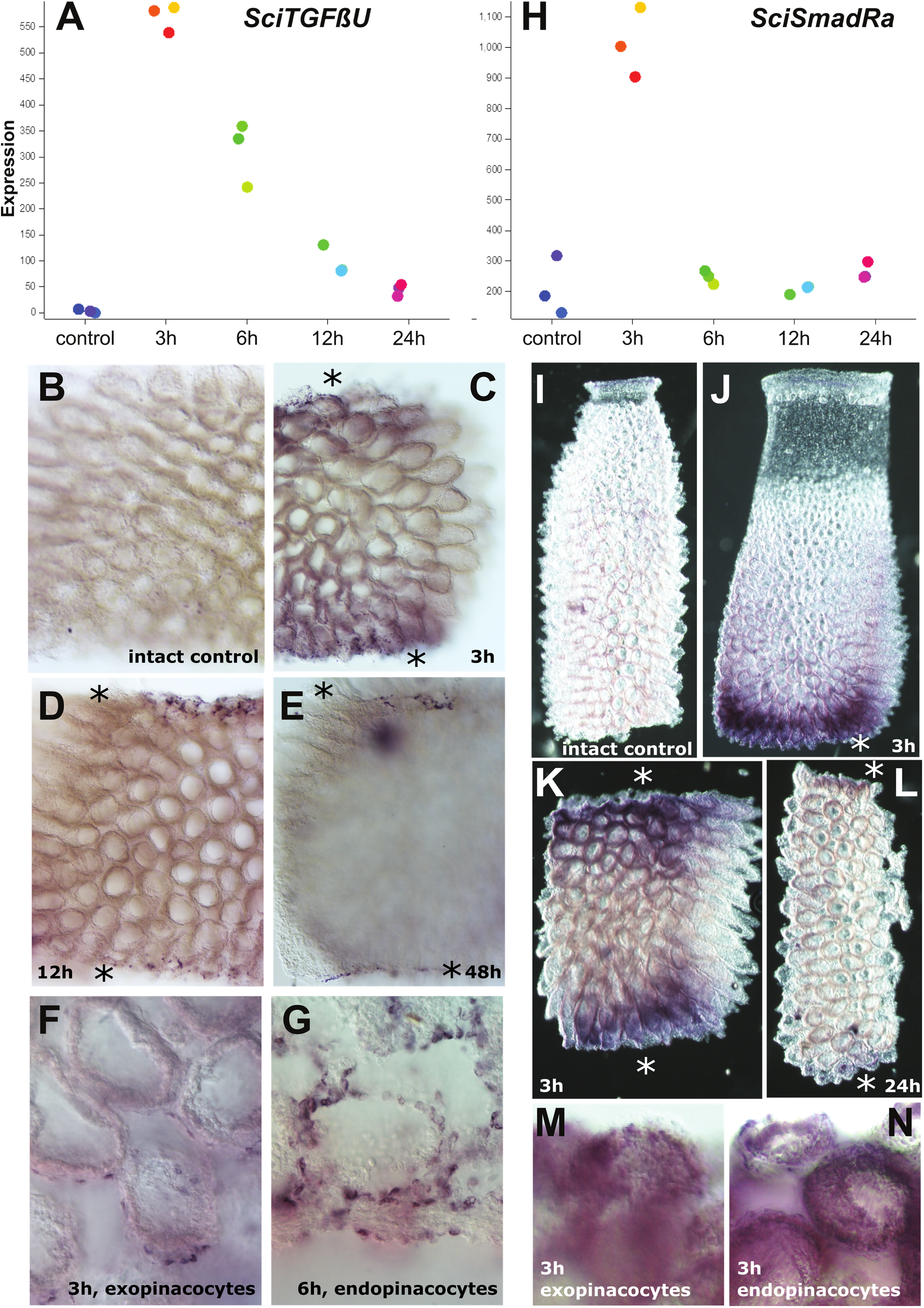
Expression profiles (top panel) and expression domains of two components of the TGFβ signalling pathway, SciTGFβU and SciSmadRa, during S. ciliatum regeneration. Gene expression levels are expressed in counts (the “expected counts” normalised for the size of the library), coloured dots represent individual replicates. Time points are indicated; the control samples were fixed before sectioning and stained in the same conditions, including time of colour development, as the experimental samples. Bottom panels show higher magnification of the regenerating regions, viewed from the outside (exopinacocytes) or the inside/atrial area (endopinacocytes). Note that to capture the gradient of SmadRa expression in regenerating samples, the staining was stopped before normal (steady-state) expression in choanocytes and mesohyl cells became visible. Asterisks (*) indicate cut surfaces.

In contrast to SciTGFβU, SciSmadRa, one of effectors of the TGFβ pathway, is expressed in choanocytes and mesohyl cells throughout the body column in non-regenerating sponges, without clear axial distribution [27]. In regenerating sponges, expression of SciSmadRa is highest (approx. 4-fold) at 3h post injury, but quickly falling to pre-injury levels by 6h based on the transcriptome analysis (Fig. 8H). *In situ* hybridisation experiments reveal that similar to SciTGFβU, SciSmadRa becomes activated within approximately three layers of radial chambers adjacent to the cut areas at the 3h time point (Fig. 8I–K). Although not apparent at the transcriptome level, and perhaps masked by relatively high steady-state levels, very weak upregulation at the injury sites could barely be visible at later stages in some samples (e.g. Fig. 8L). Strikingly, in contrast to pinacocyte-specific expression of SciTGFβU in regenerating sponges, SciSmadRa could be detected broadly across all cell types within the elevated expression domains, as evidenced by blurry rather than sharp staining pattern (Fig. 8 M, N).

## Discussion

In this study, we have explored cellular and transcriptomic dynamics of *S. ciliatum* regeneration, focusing on the wound healing stages. We firstly determined regeneration rates of transversally dissected *S. ciliatum* tissue by morphological and microscopic characterisation of distinct regeneration stages. SEM analysis revealed highly active early wound healing phases with transdifferentiation of choanocytes and collective migration of pinacocytes facilitating rapid wound closure in the absence of cell proliferation. Cell proliferation is initiated at later stages, apparently to replace cells that have been recruited to the injury site and/or undergone transdifferentiation.

*S. ciliatum* regeneration follows morphological and cellular trajectories at similar rates previously observed in other calcareous sponge species, with wound healing within 24 hours of injury and completed regeneration after 4-5 days [21, 47, 48]. The crucial morphogenetic step in early regeneration is quick wound closure by a combination of migration and spreading of endo- and exopinacoderm and transdifferentiation of exposed choanocytes to pinacocytes. Transdifferentiation is a well-known, integral mechanism of sponge regeneration across classes [17, 18, 21, 22, 47, 48]. We hypothesise that the ability of sponge cells to quickly and effectively transdifferentiate is a trait inherited from pre-metazoan ancestors, which were capable of temporal cell differentiation [47, 48, 49]. This feature remains evident in extant animal relatives, including choanoflagellates [52]. However, while a comprehensive understanding of the cellular processes in sponge wound healing has been developed, the underlying genetic basis of this process remains largely unexplored.

Here, we have generated extensive transcriptome datasets in an attempt to identify genes involved in the early stages of sponge regeneration. Integration of different RNA-seq data sets was successfully conducted using ComBat-seq [36] and allowed an intercomparison study of gene expression. Transcriptomic analyses of wound healing show that the initial process is also rapid on the transcriptome level and reveal that the early wound healing phases are partially regulated by genes with osculum-predominant or osculum-specific expression in intact sponges. This is particularly striking for the metazoan specific developmental TGFβ signalling pathway, multiple components of which are dramatically upregulated within hours of wounding. Moreover, genes associated with increased transcription, cell signalling and structural organisation are upregulated in early phases of wound healing, while genes involved in cell proliferation as well as apoptosis are downregulated. In addition, we show that although known signalling pathway components, various transcription factors and genes involved in spicule formation are differentially expressed, a significant number of uncharacterised and taxon-restricted genes may be governing wound healing in *S. ciliatum*.

Across eumetazoans, the TGFβ pathway regulates numerous cellular functions such as cell fate, apoptosis, differentiation, polarity and movement in a wide range of biological contexts including wound healing [53, 54]. This is true for a variety of structures in phylogenetically diverse systems propounding the evolutionarily ancient role of TGFβ signalling in cell behaviour during wound healing and regeneration [55, 56, 57, 58, 59, 60]. In *S. ciliatum*, the expression patterns of multiple TGFβ pathway components suggest a wide-ranging activation of cells in response to injury encompassing multiple rows of choanocyte chambers from the injury site. Interestingly, this is the same region in which increased DNA synthesis can be observed. Strikingly, one of the ligands of the pathway, TGFβU, expression of which in adult intact sponges is restricted to the apical-most cells surrounding the osculum, becomes strongly and specifically expressed in pinacocytes directly surrounding the wound area within hours of injury (Fig. 8). This expression strongly suggests involvement of the TGFβ signalling in migration of pinacocytes during initial wound closure. As the TGFβ pathway is known to be involved in collective cell migration during development, wound healing and cancer metastasis of bilaterians [61, 62, 63], our results indicate that this role of the TGFβ pathway has been conserved over 600 million years of animal evolution.

The Wnt pathway components are another major class of genes upregulated upon injury in *S. ciliatum*. While the Wnt pathway has key roles in virtually all developmental events in animals [64, 65, 66, 67, 68, 69, 70, 71], our results underscore the evolutionarily conserved role of the Wnt pathway in regeneration and axial patterning. Strikingly, transcriptomic and *in situ* hybridisation data indicate that both pathways are involved in regeneration regardless of position of the injury along the apical-basal axis of *S. ciliatum*. Similarly, expression of Wnt and TGFβ signalling pathway genes is tied to head regeneration and organiser formation in *Hydra* [56, 72, 73, 74, 75]. Intriguingly, Wnt and TGFβ pathways might also be involved in axial specification during primmorph development in reaggregation experiments of the demosponge *Halisarca dujardinii* [76].

Multiple components of the TGFβ and Wnt signalling pathways display two waves of expression following injury – the first during early wound healing (the first peak at 3-6h), and the second at the beginning of the replacement of lost structures, which we regard as the actual regeneration phase (beginning at 24h). This biphasic expression suggests that these early and late regeneration stages are indeed separate biological processes, although partly redeploying the same molecular tools. This observation supports the increasingly prevailing hypothesis in developmental biology that regeneration is at least a partial redeployment of embryonic developmental pathways [77]. A series of pairwise comparisons of differentially expressed genes revealed that a significant fraction of upregulated genes is shared between the early wound healing phase and osculum maintenance. We identified shared components of the Wnt and TGFβ signalling pathways suggesting that these developmental pathways are recruited in the regeneration of missing structures in *S. ciliatum*. While morphological and microscopic observations suggest that axial polarity is preserved during *S. ciliatum* whole body regeneration from transverse section, our current transcriptome dataset does not address the molecular mechanism of this phenomenon.

The key to the remarkable regenerative capacities of *S. ciliatum* lies in the global plasticity of differentiated cells throughout the entire body column. This remarkable feature of sponge biology is particularly emphasised by one of the most striking cell types in the animal kingdom – choanocytes, the flagella-bearing feeding cells with stem cell-like characteristics [27]. Choanocytes display a high degree of cell plasticity with the ability to transdifferentiate to endo- or exopinacocytes, which are cell types originating from a different embryological cell lineage [20]. Consistent with our observations, whole organism single cell RNA-sequencing of the demosponge *Amphimedon queenslandica* confirmed cells with transcriptional states featuring transdifferentiation intermediates between choanocytes and pinacocytes [78]. The transdifferentiation of choanocytes to pinacocytes is followed by a transition to basopinacocytes and porocytes, and thus facilitating the remodelling of a viable outer epithelial layer. Choanocytes therefore constitute the principal contributor and cell source for regeneration in species without archaeocytes (amoeboid, pluripotent cells), as *Leucosolenia sp*. (Calcarea) and *Oscarella lobularis* (Homoscleromorpha) [16, 18, 21], to which we here add the syconoid calcareous sponge *Sycon ciliatum*.

Previous research on *Leucosolenia* and *Oscarella* regeneration has demonstrated that cell proliferation was not affected by regenerative activity [18, 21]. Similarly, we observed global proliferation of choanocytes distributed throughout the choanoderm without striking differences in intact and regenerating tissue and no localisation to the wound site either at early or late stages of regeneration. This is in line with studies demonstrating that choanocytes are the most proliferative cell type in intact tissue in a number of sponge species [17, 79]. Therefore, wound healing and regeneration appear to be solely accomplished by tissue migration and transdifferentiation, traditionally termed as the “morphallactic” mode of regeneration, whereas cell proliferation probably is predominantly involved in normal tissue turnover and homeostasis at same rates in intact and regenerating tissue. Interestingly, cell kinetic analysis of early regeneration in *Halisarca caerulea* showed a decrease of proliferating choanocytes at the injury site [80]. Although no shift in cell proliferation was identified by the applied assay (EdU staining), GO enrichment analysis of the present RNA-seq data suggests a sharp drop of cell division and related GO terms such as cell cycle and DNA replication at 3-6h.

On the other hand, programmed cell death has been increasingly identified as a key prerequisite for successful wound healing and regeneration in animals in the past decade [39, 40, 79, 80, 81, 82, 83]. Only very recent studies have investigated the role of apoptosis in sponge regeneration, revealing its involvement in re-development from dissociated cells in *S. ciliatum* [28]. The importance of apoptosis for tissue remodelling remains obscure, as although two waves of apoptosis during demosponge *Aplysina cavernicola* regeneration were detected using the TUNEL assay method, no evidence for an increase of apoptosis-related genes could be found during *Halisarca caerulea* regeneration [22, 86].

In the present study, we analysed the transcriptional data for a possible role of apoptosis in calcareous sponge regeneration. We do not find significant evidence for an increase in apoptotic activity during wound healing in the first 24h after injury; in fact, there is downregulation of genes involved in the apoptotic process at 3-6h of wound healing. It is therefore possible that (at least some) sponges do not rely on apoptotic signals to induce regeneration.

As another striking observation, while we demonstrated the involvement of well-studied and expected animal developmental pathways in *S. ciliatum* regeneration, an overwhelming number of reported differentially expressed genes proved taxonomically restricted genes. Up to 20% of genes in all taxonomic groups studied so far have been identified as ‘taxon-specific’ without apparent homologs in other species [87]. Research into these genes have revealed functions in a variety of biological processes including morphogenesis and diverse regenerative mechanisms likewise in plants and animals [85, 86, 87, 88]. In the scope of this study, our investigations were focused on most significantly differentially expressed genes during wound healing as identified by the *timecourse* package. A large fraction of these genes (74 out of top 100) is unknown in function and more research will be required to uncover their putative role in sponge physiology.

## Conclusion

The present study integrates electron microscopy imaging and molecular analyses to uncover the cellular and genetic basis of *Sycon ciliatum* whole body regeneration, with focus on wound healing. We report a highly dynamic wound healing phase involving major cellular events such as concerted migration and transdifferentiation of epithelial cells, potentially governed by ancient animal developmental signalling pathways as well as numerous taxonomically restricted genes. This first transcriptomic characterisation of calcareous sponge tissue remodelling expands the number and phylogenetic range of available models for animal regeneration and will be useful in our endeavour to better understand the evolution of one of the most remarkable animal traits.

## Methods

### Sample collection and experimental procedures

Adult *Sycon ciliatum* specimens were collected from Norwegian fjords near Bergen in October, outside the reproductive season. All samples for RNA-Seq were collected using pools of 8-10 sponges per replicate, with three replicates per experiment.

#### Maintenance of body plan in intact sponges

Samples were constructed from axially dissected specimens as described in [27]. Briefly, intact sponges were cut to obtain top (osculum), bottom (basal part), and two adjacent middle sections: mid1 being closer to the top and mid2 to the base.

#### Wound healing

Sections of the body column were cut and kept in Petri dishes with seawater (excluding apical and basal parts). RNA samples were taken at 3, 6, 12 and 24 hours post sectioning to capture early regeneration processes. At the same time, intact sponges were kept in dishes in the same way and middle sections were collected at the same time as 3 and 6 hours experimental samples (combined control 1) and 12 and 24 hours experimental samples (control 2) to account for the effect of keeping sponges under laboratory conditions (as opposed to field collection).

### Light and electron microscopy

For microscopic investigation, samples were fixed in 2.5% glutaraldehyde (Ted Pella, Redding), four volumes of 0.2 M cacodylate buffer and five volumes of seawater (1120 mOsm) for 2 h and post-fixed in 2% OsO_4_ (Spi Supplies, West Chester) in seawater at room temperature for 2 h. After fixation, samples were washed in 0.2 M cacodylate buffer. After postfixation, specimens were subjected to spicule dissolution in 5% ethylenediaminetetraacetic acid (EDTA) (Sigma-Aldrich, St. Louis) solution for 2 hours at RT. Finally, specimens were dehydrated in an ethanol series at RT and stored in 70% ethanol at 4°C and were embedded in Araldite resin (Sigma-Aldrich). Semithin (1 μm) and ultrathin sections (60 – 80 nm) were cut on an Ultramicrotome PowerTome XL (RMC Boecke-ler, Tucson), and then stained with 1% toluidine blue. The semithin sections were studied under a Leica DM5000B (Leica). Digital photos were taken with a Leica DMLB microscope (Leica) using the Evolution LC color photo capture system (Media Cybernetics, Rockville). Ultrathin sections were contrasted with uranyl acetate and lead citrate, and observed under a Jeol JEM-1400 transmission electron microscope (TEM). For scanning electron microscopy (SEM), fixed specimens were critical-point-dried, sputter-coated with gold-palladium, and observed under a Hitachi S 570 SEM.

### Cell proliferation assay

Proliferating activity was detected by incorporation of labeled thymidine analog, 5-ethynyl-2′-deoxyuridine (or EdU), during S-phase of cell cycle, with subsequent visualisation of labeled DNA by click-reaction [91]. EdU and the reactive Alexa Fluor™ 488 dye used according the manufacturer instructions to Click-iT™ EdU Imaging Kit (cat.no. C10086; Invitrogen, USA). Five regenerates (cross section of the body) at each timepoint – 3, 24, 48, 72 and 96 h – were studied. Unwounded sponges used as positive control; equivalent amount of DMSO without EdU was added to seawater in negative control samples to verify nonspecific background. Regenerates placed in filtered sea water supplied with 300 μM EdU, diluted from 250 mM stock in DMSO. Specimens were placed in EdU 1 h before reaching the exact timepoint, incubated 1 h at 16°C in Petri dishes, and fixed in 4% PFA in 0.1M PBS (pH 7.4) overnight. Fixed specimens were washed in 0.1M PBS with addition of 0.1% Tween-20 three times (10 min each), permeabilised with 0.5% Triton X-100 in 0.1M PBS during 15 min on a shaker, and blocked in 1% bovine serum albumin in PBS. After rinsing in PBS, samples were incubated in staining mix during 1 h at room temperature in the dark. Then colonies were rinsed 0.1M PBS with addition of 0.1% Tween-20 two times (10 min each). Nuclei were counterstained with DAPI at 1 μg/ml. Specimens were mounted in glycerol with DABCO addition as an antifade and examined using the confocal laser scanning microscope Leica TCS SP5. The images were processed by ImageJ software (FiJi).

### Whole mount *in situ* hybridisation

*In situ* was performed as previously reported in [92].

### RNA sequencing and transcriptome assembly

RNA was extracted using the AllPrep DNA/RNA Mini Kit (Qiagen) according to manufacturer’s protocol. Complementary DNA and genomic libraries were constructed with the TruSeq RNA Library Prep Kit v2 (Illumina) and sequenced using Illumina technology at The Norwegian High-Throughput Sequencing Centre. After demultiplexing pooled samples and read trimming using *trimmomatic v0.38*, sequences were quality controlled with FastQC [93, 94]. Transcriptome was assembled *de novo* from all the RNA-Seq libraries using Trinity pipeline with default parameters [95]. Transcripts assembled by Trinity were clustered with cd-hit at 95% [96] identity and the longest transcript from each cluster was selected. Selected transcripts were then cleaned from alien (non-sponge) contaminations by aligning on them (with the blat tool [97]) the ‘clean’ Illumina reads from sequencing of genomic library built from *S. ciliatum* larvae collected and washed in filtered sea water. Only transcripts targeted by at least two ‘clean’ reads were selected for further analyses. Open reading frames were predicted with transdecoder [98] using evidence from protein domain prediction (hmmscan [99] against Pfam A database [100]).

### Gene expression analysis

Gene expression values were calculated as the “expected counts” using *RSEM v1.3.1 (rsem-calculate-expression)* as a wrapper for *Bowtie2* by mapping the reads to the *S. ciliatum* transcriptome and formatted into a data matrix (*rsem-generate-data-matrix*) [101, 102]. To integrate intact sponge and wound healing experiments, data was corrected for batch effects using *ComBat-seq* [36]. Differential gene expression (DEG) and principal component (PCA) analyses were performed using the *DEseq2* package available in *R* [103, 104]. Replicate naming (r1-r3) refers to samples within an individual experiment. For DEG analysis, *apegalm* shrinkage of log2 fold change estimates (*lfcShrink*) was used [105]. Genes with a log2 fold change > ±1 (2-fold or greater) and adjusted p-value < 0.05 were considered significantly differentially expressed. Counts were transformed for PCA applying a regularised log transformation to normalised counts in *DEseq2*. DEG’s in wound healing time-course data were identified using the *timecourse* package, which applies a multivariate empirical Bayes analysis approach [41, 42]. As this package was developed for microarray data, counts were *voom* transformed beforehand [106, 107]. Gene ranking is based on Hotelling T^2^-statistics in the order of evidence of nonzero means incorporating the correlation structure of time points, replication and moderation. Gene clustering by expression profiles was performed with *clust* using default parameters and adjusted normalisation type (*normalisation code 101 31 4*) [37]. Heat maps to visualise gene expression patterns were built using the *pheatmap* package in *R* and the package *GOseq* was used for gene ontology analysis [38].

## Acknowledgements

This work was supported by Sars International Centre for Marine Molecular Biology (Bergen, Norway) and Australian Research Council through ARC Centre for Excellence for Coral Reef Studies (CE140100020) and Future Fellowship (FT160100068) funding. Sequencing was carried out at the Norwegian High-Throughput Sequencing Centre funded by the Norwegian Research Council.

## Author contributions

Conceived and designed experiments: Maj.A., Mar.A.; developed protocols and reagents: Mar.A., G.A., I.B.; performed experiments and collected data: M.L., S.L., D.T., A.E., Maj.A; performed analyses: C.C., Mar.A., Maj.A.; wrote and edited the manuscript: C.C., A.E., Maj.A., Mar.A., D.P., I.B.

